# Inoculation by mosquito induces durable differences in serological profile in non-human primates infected with DENV1

**DOI:** 10.1101/2020.09.28.315218

**Authors:** Jayant V. Rajan, Michael McCracken, Caleigh Mandel-Brehm, Greg Gromowski, Simon Pollett, Richard Jarman, Joseph L. DeRisi

## Abstract

Natural dengue virus (DENV) infections are delivered by mosquito bite but how the route of inoculation route could shape the humoral immune response is not well understood. Here, we serologically profiled 20 non-human primates (NHP) from a prior study of DENV1 infection in which the animals were inoculated by mosquito (N=10) or subcutaneous injection (N=10). Using a comprehensive, densely tiled and highly redundant pan-flavivirus programmable phage library containing 91,562 overlapping 62 amino acid peptides, we produced a high-resolution map of linear peptide sequences enriched during DENV seroconversion. We found that serological profiles in mosquito-inoculated and subcutaneously-inoculated animals were similar up to 90 days after primary infection, but diverged at 1 year. We found differences in sero-reactivity, as indicated by the median area under the curve (AUC) in the Envelope (E; residues 215-406; p < 0.08), and Nonstructural-3 (NS3; residues 549-615; p < 0.05) proteins in mosquito-inoculated versus subcutaneously-inoculated animals. Within the E protein, residues 339-384 in domain III accounted for >99% of the total AUC difference across residues 215-406. Antibody breadth did not vary by mode of inoculation. The differential reactivity to E domain III (EDIII) seen by phage display validated orthogonally by ELISA, but did not correlate with late neutralization titers. Serological profiling of humoral immune responses to DENV infection in NHP by programmable phage display demonstrated durable differences in sero-reactivity by route of inoculation. These findings could have implications for DENV diagnostics and evaluation of vaccines.

**IMPORTANCE:** Dengue virus (DENV) infections are transmitted by mosquito bite, but how being infected by mosquito bite affects the immune response is not known. In this study, we analyzed antibodies produced by rhesus macaques infected with DENV using programmable phage display, a high-throughput method for characterizing what viral protein derived peptides serum antibodies bind to. We found that while there was no difference in antibody binding profiles at early timepoints post-infection, at one year post-infection, there were substantial differences in the antibody binding profiles of macaques who were infected by mosquito bite versus those that were infected by injection. In general, antibodies in the macaques inoculated by mosquito maintained higher levels of sero-reactivity, with a strong signal still present one year post-infection. The findings we report could have implications for DENV diagnostics and evaluation of DENV vaccines.

## INTRODUCTION

Dengue virus (DENV) is an arbovirus that has been a major cause of global infectious disease morbidity and mortality for decades(1,2). Clinical manifestations of DENV infection can range from asymptomatic disease to severe sepsis with coagulopathy, with many infected persons developing symptomatic, non-severe disease, characterized by fever, rash and arthralgias (3,4). Severe disease occurs primarily in the setting of heterotypic serial infections with any combination of two of the four DENV serotypes and is more common in children than adults (5).

Primary DENV infection in humans generates type-specific neutralizing antibodies to the infecting serotype as well as transient, cross-reactive neutralizing antibodies against other DENV serotypes (6,7). Secondary infection has a crucial role as it elicits type-specific antibodies to the second serotype and, more importantly, durable, cross-reactive, broadly neutralizing antibodies (8). Much of the current understanding of DENV immunity in humans derives from studies of human sera as well as studies of monoclonal antibodies isolated from B-cells of persons infected with DENV. Such studies have focused on the analysis of natural DENV infections delivered by *Aedes* mosquitoes.

There are currently multiple tetravalent DENV vaccines that have completed or are in phase III clinical trials, each of which seeks to recapitulate the durable immunity that results from authentic serial heterotypic DENV infections (9–11). While the vaccination strategies differ between the advanced vaccine candidates, all are delivered by subcutaneous or intramuscular injection, a very different mode of delivery than inoculation by mosquito.

Non-human primate models of DENV infection are the best animal approximation of virus replication in humans. In this study, programmable phage display immunoprecipitation sequencing (PhIP-Seq), a high throughout serological profiling technique was used to characterize serological responses to primary DENV1 infection as well as both homologous (DENV1) and heterologous (DENV2) challenge in rhesus macaques infected by mosquito bite and by subcutaneous injection (12–14). A densely tiled, redundant set of peptides, encompassing flavivirus sequence variability as represented by publicly available sequence data was designed to allow for detailed mapping of both the homotypic and heterotypic response to DENV-derived peptides. These data are generated in a single, high throughput assay that requires minimal serum volume, in contrast to standard serum depletion and neutralization assays. The results of this study suggest that the route of infection can have a substantial effect on the immune profile to peptide antigens, with viral inoculation by mosquito inducing a different, durable pattern of sero-reactivity as compared to subcutaneous injection. These results could have implications for DENV diagnostics, sero-epidemiology and current DENV vaccination strategies.

## RESULTS

### Inoculation of NHP with DENV1 by mosquito or by subcutaneous injection induces humoral responses of similar magnitude with similar kinetics

The programmable phage library used here consists of 91,562 overlapping 62 amino acid peptides derived from 74 flaviviruses (see Methods). Next Generation Sequencing (NGS) of the packaged phage display library confirmed > 96% representation of designed peptides, with 96% of them present at ≤ 1 read per 100,000 reads (Supplemental Figure 1). The experimental procedure for immunoprecipitations is depicted in figure 1A. For the 10 non-human primates (NHP) inoculated by mosquito, the number of enriched peptides relative to the pre-infection baseline sample at 7, 35, 90 and 365 days post-infection (p.i.) ranged from 22-344, 111-416, 22-540, and 28-501. For the 10 NHP inoculated by subcutaneous injection of DENV1 the number of enriched peptides at 7, 35, 90 and 365 days p.i. ranged from 26-93, 88-930, 64-789 and 31-786.

**Figure 1.**
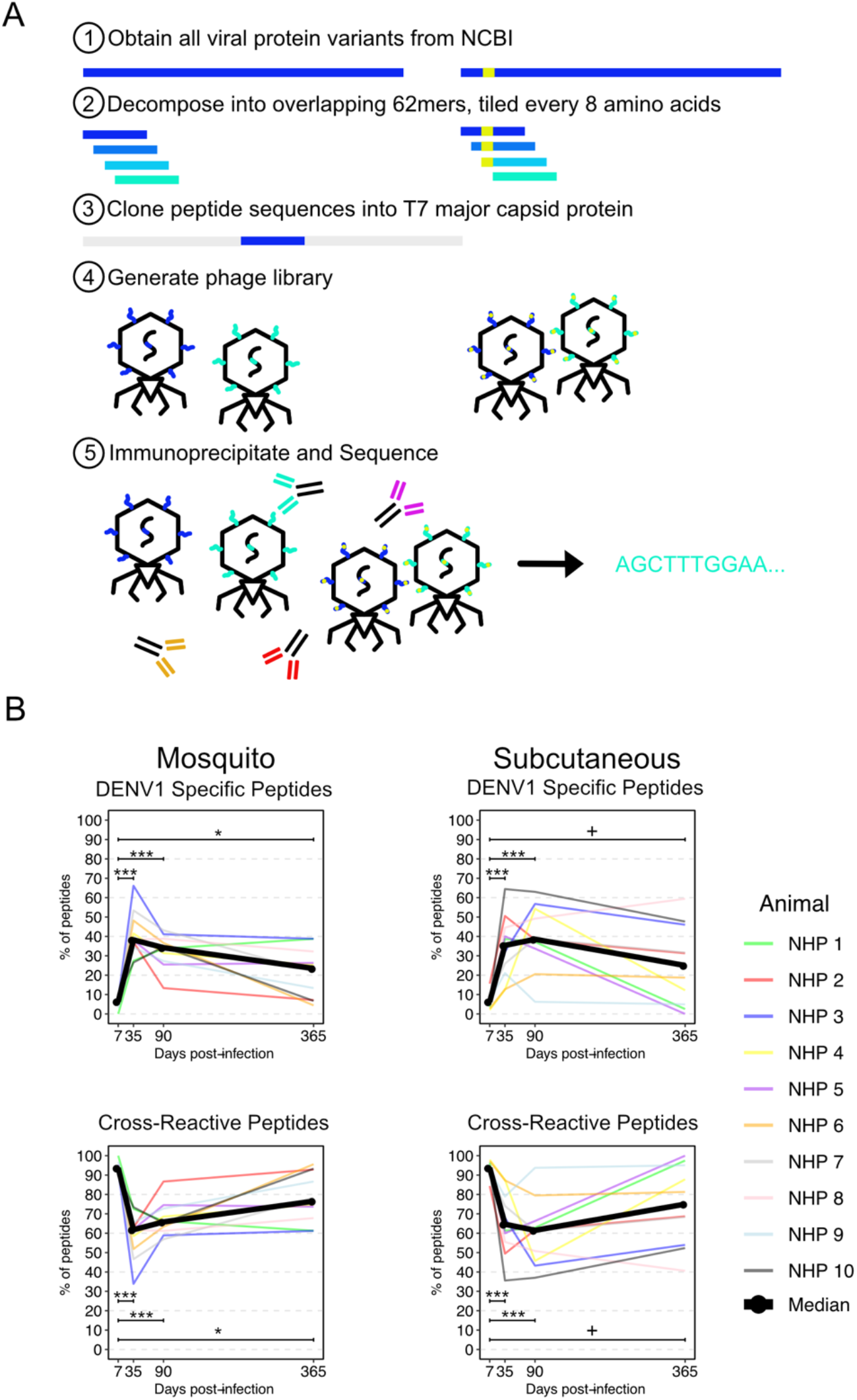
Phage display measurement of total sero-reactivity to dengue virus is similar in nonhuman primates infected by mosquito and subcutaneous infection. A redundant, densely tiled set of peptides encompassing 74 flaviviruses reported to infect humans was designed, validated and used for phage immunoprecipitation sequencing (B). Animal to animal signal variability was high in both the mosquito-infected (N=10 animals) and subcutaneously infected (N=10 animals) groups. DENV1 specific median sero-reactivity (>90% similar to DENV1 only) in both groups was increased at 35, 90 and 365 days post-infection (*** p < 0.001; * p < 0.05; + p < 0.1) compared to 7 days post-infection. Cross-reactive median sero-reactivity declined in both groups as DENV1-specific sero-reactivity increased.

The proportion of DENV1-specific peptides, defined as peptides that were ≥ 90% similar to DENV1 but not to any other DENV serotype, peaked at 35 days p.i. and decreased at 90 and 365 days p.i. (Figure 1B, left). The difference in the median proportions of DENV1-specific peptides for each of these timepoints as compared to 7 days p.i. was statistically significant (Mann-Whitney U test) and this was true regardless of the route of inoculation. At 7, 35, 90 and 365 days p.i., the median proportions of DENV1-specific peptides in the mosquito-inoculated group were 6.4% [interquartile range (IQR) 4.6-7.8%], 38.2% [IQR 34.9-66.1%; p < 0.001], 34.2% [IQR 28.3-43.1%, p < 0.001], and 23.4% [IQR 8.7-38.8%, p < 0.05] in the mosquito-infected group. For the subcutaneously inoculated group, the median proportions at days 7, 35, 90 and 365 p.i. were 6.3% [IQR 3.9-7.6%], 35.4% [IQR 22.3-43.4%, p < 0.001], 38.5% [IQR 34.6-53.0%, p < 0.1].

Cross-reactive peptides, defined as peptides that were ≥ 90% similar to DENV1 as well as at least one other DENV sero-type or as peptides that were <90% similar to DENV1, showed a different pattern, decreasing from day 7 p.i. as the proportion of DENV1-specific antibodies increased. As with DENV1-specific peptides, this pattern did not differ by route of inoculation. In the mosquito-inoculated group, the median proportion of cross-reactive peptides at 7, 35, 90 and 365 days p.i. were 93.6% [IQR 92.2-95.3%], 61.8% [IQR 53.5-65.1%; p < 0.001], 65.8% [IQR 61.8-71.7%; p < 0.001], 76.5% [IQR 69.3-95.6%; p < 0.05]. In the animals inoculated by subcutaneous injection, the median proportions of cross-reactive peptides were 93.7% [IQR 92.4-96.1%], 64.6% [IQR 56.6-77.7%; p < 0.001], 61.5% [IQR 47.0-65.4%; p < 0.001] and 75.0% [IQR 57.8-93.2%; p < 0.1].

### Primary infection by mosquito induces durable differences in serological profile compared to subcutaneous inoculation

Minimal sero-reactivity was observed at 7 days post-infection, but by 35 days post-infection, the median antibody coverage for each group of 10 animals enriched for 17.3% (mosquito) and 14.3% (subcutaneous) of the DENV1 proteome respectively (Figure 2). This coverage focused on the envelope (E), nonstructural-1 (NS1), nonstructural-3 (NS3) and nonstructural-5 (NS5) proteins. In the NHP inoculated by mosquito, the E, NS1, NS3 and NS5 proteins represented 44.2%, 20.8%, 17.9%, and 15.1% of the enriched regions. For NHP inoculated subcutaneously the same four proteins represented 48.0%, 19.6%,13.6% and 9.2% of the targeted regions.

**Figure 2.**
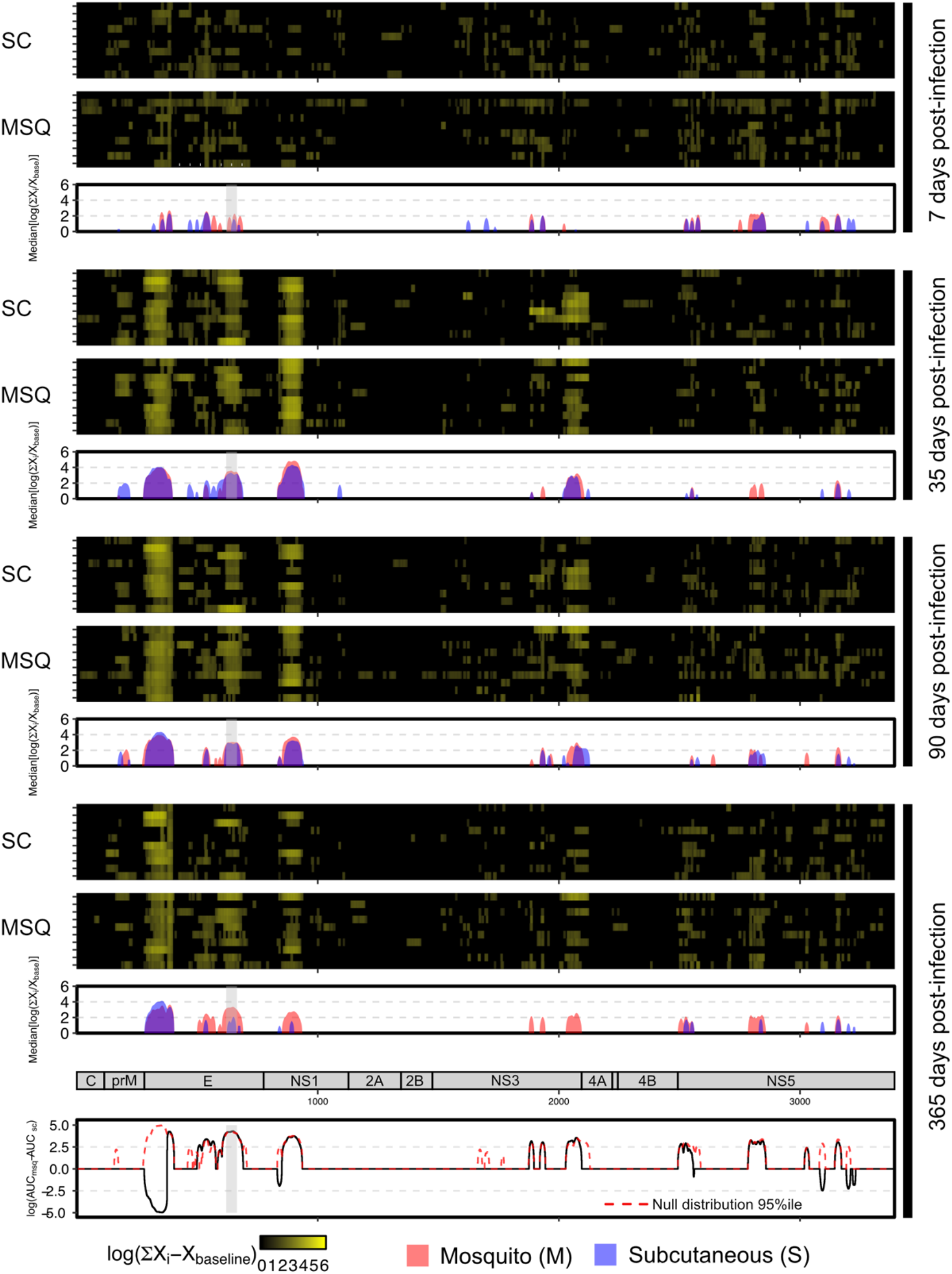
Non-human primates infected by mosquito demonstrated persistent differences in sero-reactivity across the DENV1 proteome. Significantly enriched peptides were aligned to the DENV1 reference proteome and median enrichments computed. The E, NS1, NS3 and NS5 proteins were the most frequently targeted and by 365 days post-infection, while reactivity to E, NS1 and NS3 remained high in subjects infected by mosquito (red), they declined in those infected by subcutaneous injection (blue). Within the E protein, the differential reactivity centered on amino-acid residues 339-384 (grey rectangle), with a statistically significant difference in the AUC. The 95%ile of the null distribution of the AUC difference is shown with a red dotted line.

At 90 days post-infection, the median antibody coverage enriched for 16.3% and 15.3% of the DENV1 proteome in NHP inoculated by mosquito and subcutaneous injection. Of the enriched regions, the E, NS1, NS3 and NS5 proteins represented 43.0%, 15.5%, 18.1%, and 16.1% in mosquito-inoculated NHP versus 36.6%, 17.0%, 15.6% and 21.2% in subcutaneously inoculated NHP.

At 365 days post-infection, the median antibody coverage targeted 16.8% and 7.4% of the DENV1 proteome in NHP inoculated by mosquito and subcutaneous routes. Of the enriched regions, the E, NS1, NS3 and NS5 proteins represented 46.5%, 13.4%, 14.6%, and 25.5% in mosquito-inoculated NHP versus 62.1%, 8.3%, 0% and 29.6% in subcutaneously inoculated NHP. The area under the curve (AUC) was calculated as a measure of antibody breadth and magnitude and showed distinct patterns at this late timepoint by route of inoculation (Figure 2, bottom). We observed differences in the AUC spanning residues 549-615 of the NS3 protein with a larger AUC in the mosquito-inoculated group (log_10_[AUC_msq_ - AUC_sc_] = 4.03, p < 0.05). We obtained similar results in the E protein, with a region spanning residues 1-91 (domain I) having a larger AUC in the subcutaneously-inoculated group (log_10_[AUC_sc_ - AUC_msq_] = 5.62, p < 0.075) and another region spanning residues 215-406 (domain III) where the AUC was larger in the mosquito-infected group (log_10_[AUC_msq_ - AUC_sc_] = 5.02, p < 0.075). For the latter region, 97.6% of this AUC difference (log_10_[AUC_msq_ – AUC_sc_] = 4.90, p < 0.1) was accounted for by residues 339-384 in domain III. While this region was present in both mosquito- and subcutaneously infected groups at 35 and 90 days post-infection, by 365 days post-infection, as indicated by AUC difference, it had waned substantially in the subcutaneously infected group.

### Persistence of antibodies targeting EDIII after primary infection with DENV1 by mosquito but not subcutaneous injection validated by ELISA

We cloned the segment of the E protein mapping to ED III into an expression vector and used the overexpressed protein uas the antigen in an ELISA. OD450_Timepoint_/OD450_baseline_ for days 7, 35, 90, and 365 post infection for each animal in both groups was concordant with PhIP-Seq results (Fig 3A). For the mosquito group, the median OD450_Timepoint_/OD450_baseline_ ratio for 35 (1.50, p=0.0004), 90 (1.35 p=0.0004) and 365 (1.41, p=0.03) days post primary infection showed statistically significant increases (147%, 132% and 138%, respectively) relative to the median ratio for 7 days post primary infection (1.02). For NHP infected by the subcutaneous route, the median OD450_Day post infection_/OD450_baseline_ ratios for days 35 (1.53, p=0.001) and 90 (1.17, p=0.003) post primary infection also showed statistically significant increases (146% and 112%, respectively) relative to the median day 7 post primary infection ratio (1.04), but the median ratio 365 days post primary infection (1.05) did not. The median OD450_Day post infection_/OD450_baseline_ ratio for the mosquito group at 365 days post primary infection also showed a statistically significant increase (145%) relative to the ratio 365 days post primary infection for the subcutaneous group (p < 0.05).

**Figure 3.**
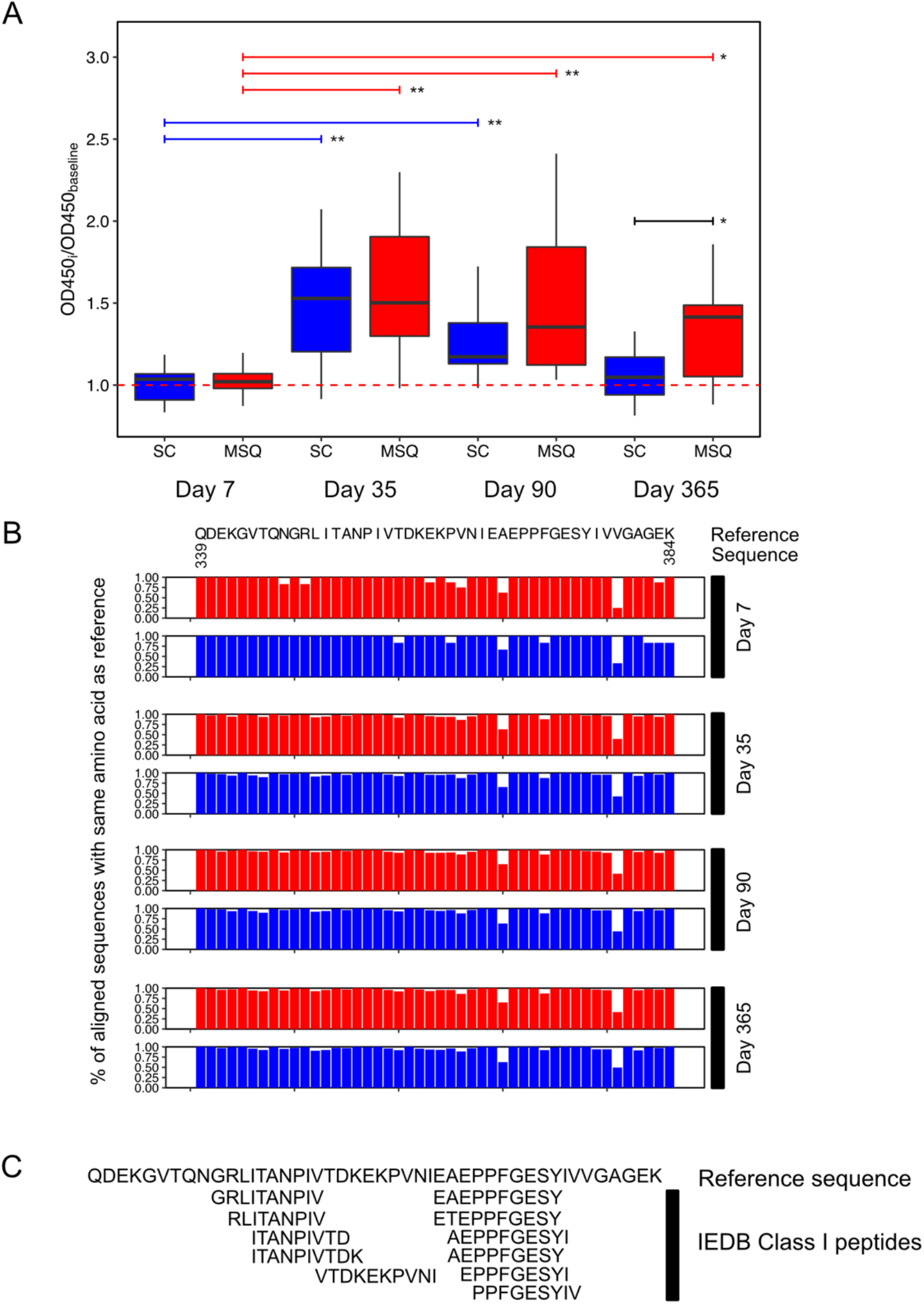
Persistent differences in EDIII sero-reactivity validate by ELISA with PhIP-Seq enriched peptides to this region showing no differences in antibody breadth by the route of inoculation. The region of differential reactivity was cloned into a mammalian expression vector, overexpressed in HEK293T cells and used as the antigen in an ELISA. By ELISA, EDIII seroreactivity in the mosquito-infected (red) but not in the subcutaneously infected (blue) group was significantly elevated at 1 year post-infection as compared to 7 days post-infection (A; ** p < 0.01; * p < 0.05). Antibody breadth was similar in both groups, with only 2 positions (E_368_, E_379_) with < 70% conservation of the reference sequence residue (B). Several MHC Class I peptides, but no Class II epitopes from the immune epitope database (IEDB) mapped to the region of interest (C).

### Route of inoculation does not affect breadth of antibodies targeting ED III

Multiple sequence alignment of the peptide sequences enriched in mosquito-inoculated and subcutaneously inoculated NHP were done against the DENV2 reference sequence (Fig 3B). At 7 days p.i. at 38/46 of the positions in the targeted region (E_339-384_), more than 90% of the aligned peptides covering each position were identical to the reference sequence regardless of the route of inoculation. At both 35 and 90 days p.i., 42/46 positions were > 90% conserved in the mosquito group and 41/46 in the subcutaneous group. By 365 days p.i., the proportions of >90% conserved residues were 43/46 and 42/46 in the mosquito and subcutaneously-inoculated groups. In both groups, only 2 positions, Alanine_368_ and Valine_379_ demonstrated <70% sequence conservation relative to the reference sequence.

### MHC Class I epitopes map to the region of differential sero-reactivity in EDIII

The immune epitope database (IEDB) was queried for validated MHC Class I, MHC Class II and B-cell linear epitopes contained in the differentially sero-reactive region in EDIII. A total of 11 MHC Class I epitopes were found to map to this region, but no MHC Class II or B-cell linear epitopes (Fig 3C).

### EDIII targeting did not correlate with late neutralization titer

The ELISA data from 365 days p.i. did not correlate with late (13 month) neutralization titers for either the mosquito-inoculated or the subcutaneously inoculated group, with Pearson correlation coefficients of −0.34 and 0.34 for each group, respectively. We next determined whether the phage display results for the targeted region of EDIII correlated with the ELISA data, calculating the Pearson correlation between the sum of the median enrichments spanning residues 339-384 and the ELISA OD 450 ratios. The correlation coefficients were 0.84 and 0.52 for the mosquito and subcutaneously inoculated NHP. Because the ELISA data for the mosquito group correlated well with phage display results, we next examined correlations between the summed enrichments of all 45 amino acid windows across the DENV1 proteome for the 365 days p.i. profiles and the 13 month neutralization titers (Fig 4). The maximum correlation we observed in the mosquito-infected group was 0.91 for a 45 amino-acid window centered at amino acid residue 2618 in the NS5 protein, although several other amino acid windows also achieved a correlation of ≥ 0.9, predominantly in the NS3 and NS5 proteins. At this same timepoint, the maximum correlation observed for any 45 amino acid window and the 13 month neutralization titers in the subcutaneously infected group was 0.62.

**Figure 4.**
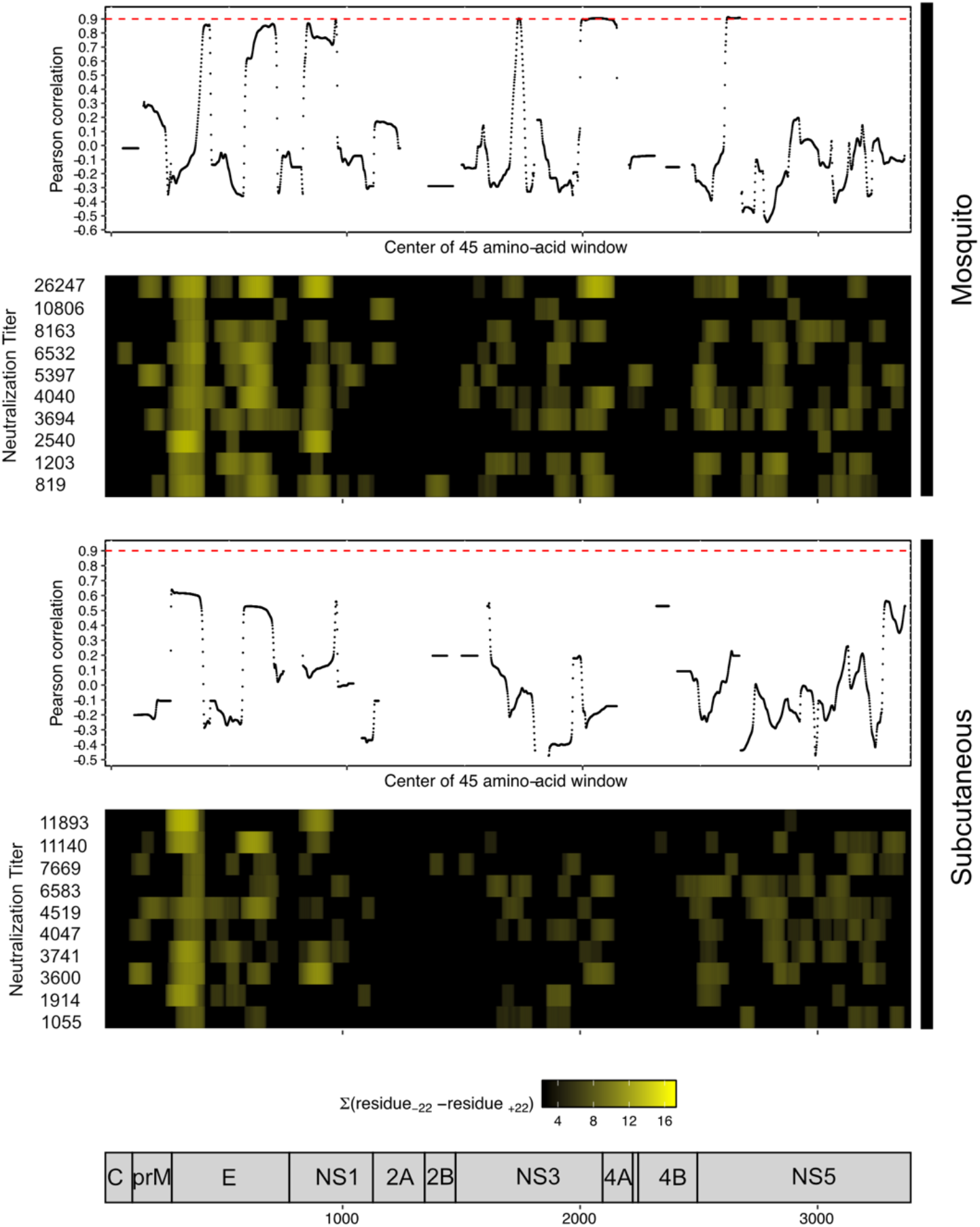
Phage display enrichments in NS3 and NS5 proteins correlate with late neutralization titers. Enrichment data from PhIP-Seq experiments 365 days post-infection was correlated with 13 month neutralization titers, by calculating the summation of enrichments for all 45 amino-acid windows across the DENV1 proteome (shown in heat maps). No 45-amino-acid windows in the E protein achieved a Pearson correlation of greater than 0.9 in either group (red dashed line). However, in the mosquito-infected group several windows in the NS3 and NS5 proteins did have Pearson correlation coefficients of ≥ 0.9. Neutralization titers from each animal (geometric mean titer) are shown on the left.

## DISCUSSION

The primary goal of this work was the determination of a high-resolution map of seroreactivity after DENV infection in non-human primates using a comprehensive set of tiled linear 62 amino acid peptides displayed on phage. Secondarily, we sought to determine whether the route of inoculation resulted in differences in this map. While serial heterotypic DENV infection by mosquito bite induces long-term immunity, it is unknown whether this route of inoculation affects the humoral immune response is unknown. In this study, serological profiling of primary DENV1 infections by programmable phage display demonstrated statistically significant differences in immune profiles in NHP inoculated by mosquito bite versus subcutaneous injection. Sero-reactivity to the DENV E, NS1, NS3 and NS5 proteins was observed at 35 and 90 days post-primary infection and did not vary by route. At one year post-infection, however, reactivity to the E and NS3 proteins demonstrated statistically significant increases in mosquito-inoculated animals versus subcutaneously-inoculated animals. The sustained differential reactivity to the E protein centered on residues 339-384 in domain III (ED III), a result that orthogonally validated by ELISA. There was no difference in antibody breadth overall, as measured by sequence diversity, by route of inoculation. Although none of the reported validated MHC Class II or B-cell linear epitopes mapped to the targeted EDIII region, multiple validated MHC Class I epitopes did. ELISA results for the targeted ED III region did not correlate with 13-month neutralization titers, but further analysis of phage data suggested several regions in the NS3 and NS5 proteins that did do so, but only in animals infected by mosquito.

There are multiple potential causes for the late differences in sero-reactivity observed after mosquito and subcutaneous infection with DENV we observed. Mosquito saliva, introduced during biting, increases DENV infectivity and enhances viral replication and could have resulted in a higher more immunogenic inoculum (16–18). Arguing against this possibility, however, is that there was a delayed onset of viremia in mosquito-inoculated animals (14). Alternatively, the immunomodulatory effects of mosquito saliva, including increased recruitment of inflammatory cells (including dendritic cells), downregulation of IFN-g and TNF-a, and upregulation of IL-4 and IL-10, suggest its ability to function as an adjuvant role in DENV infection (19–22). Given the large number of uncharacterized proteins in mosquito saliva, it is certainly plausible that one or more of these could be an adjuvant (23–25).

The virus stock used to infect NHP in the parent study was produced in Vero cells and is another potential explanation of our findings. For mosquito infections, it was injected into the thoracic cavity of mosquitoes which were then used to infect NHP 14 days later, allowing for multiple viral replication cycles in mosquitoes. Post-translational modifications (PTM) of the DENV E protein, which contains two well-documented glycosylation sites, play an important role in viral attachment and entry (26,27). PTM machinery in mammalian and insect cells is different, with mammalian cells capable of producing more complex structures (28). These PTM differences do not affect viral infectivity, but could affect the innate immune response and thus shape the adaptive immune response to DENV infection (29). Even so, mosquito-derived virus would only be present for early rounds of infection, making this an unlikely cause of the differences observed.

The DENV E protein is the primary target of the neutralizing antibodies (nAb) which are crucial for immunity to DENV, with ED III mediating viral attachment through its lateral ridge (30). Anti-EDIII antibodies, however, constitute only a small proportion (< 10%) of serum DENV nAb in humans and are not required for DENV neutralization (6,31,32). It is thus unsurprising that we observed no correlation between ED III ELISA results and 13-month neutralization titers, although our phage display results do suggest sero-reactivity to parts of NS3 and NS5 as candidate biomarkers. The functional significance of the differential sero-reactivity to EDIII we observe is unclear, but at least two DENV vaccine do appear to target this region and interest remains in EDIII-based vaccination strategies (33,34)(35–37). Finally, independent of its role in neutralization, the fact that the region we identify contains multiple reported Class I epitopes does support its immunological relevance. While it remains surprising that we identified no MHC Class II epitopes, this may be due to biased database representation, with roughly three times more Class I epitopes (1195) as Class II epitopes (428).

Using linear peptides displayed on bacteriophage to understand humoral immune responses has some limitations. DENV nAbs are thought to be against conformational epitopes defined by non-contiguous residues in the E protein and cannot be represented in 62 amino acid peptides (38,39). Phage displayed peptides also lack eukaryotic PTM that could be important for antibody recognition. Nevertheless, using this approach to characterize the full spectrum of DENV sero-reactivity in the context of natural infection can provide valuable insights about DENV immunity. Finally, while NHP are the closest model to approximate DENV replication in humans, the findings we report here represent a single group of animals and need to be verified to determine if they generalize.

There are three DENV vaccine candidates currently that have either completed or are in the midst of phase III clinical trials and serologic diagnostics for DENV remain challenging. The results reported here show that in an NHP infection model, infection by mosquito can result in very different humoral immune profiles than infection by subcutaneous injection and that these are long-lasting. While the precise causes for these differences and their functional consequences remain to be determined, the findings reported here suggest that natural immunity to DENV infection is influenced by vector delivery and should be investigated further.

## METHODS

### Dengue virus cell culture and non-human primate infections

Culture of the DENV1 and DENV2 strains and NHP infections have previously been described (14).

### Phage display library design and construction

All protein sequences present in the National Center for Biotechnology Information (NCBI) GenBank database as of November 4, 2017 were downloaded for 74 flaviviruses known to infect humans. For each virus a multiple sequence alignment was performed and the aligned sets of sequences divided into sets of overlapping 62 amino acid peptides. Consecutive peptides overlapped by 54 amino acids. The full set of peptides was collapsed on 98% sequence similarity, resulting in a final set of 91,562 flavivirus-specific peptides. Nucleic acid sequences coding for each peptide were designed using randomized sets of codons such that overlapping regions of consecutive peptides did not have the same nucleic acid sequence. Common 5’ (GCAGGAGTAGCTGGTGTTGTG, coding for AGVAGVV) and 3’ (TGATAAGCATATGCCATGGCCTC) linker sequences were appended to each oligonucleotide coding sequence, all sequenced outputted to a FASTA format file and sent to Agilent, Inc. for synthesis. All bioinformatic analysis was done using R and R studio.

Lyophilized oligonucleotides were resuspended in 10 mM Tris-HCl-1mM EDTA, pH 8.0 at 2 nM. The oligonucleotide stock was diluted to 0.2 nM and used as the template for PCR. The forward primer was TAGTTAAGCGGAATTCAGCAGGAGTAGCTGGTGTTGTG and the reverse primer was ATCCTGAGCTAAGCTTGAGGCCATGGCATATGCTTATCA. Cycling conditions used were: 98C x 30s followed by 20 cycles of 98C x 5s, 70C x 15s and 72C x 10s and terminating with 72C x 2”. A total of 4 independent PCR reactions (50ul reactions) were pooled together and cleaned using Ampure XP beads at 1X. Bead clean-up followed the manufacturer’s protocol. Bead-cleaned PCR products were quantified using Qubit reagent. One microgram (1ug) of PCR was digested with 1ul of EcoR I-HF (New England Biolabs) and 1ul Hind III-HF (New England Biolabs) in 1X Cutsmart buffer and a final reaction volume of 50ul for 1H at 37C. Restriction digests were heat inactivated for 10” at 65C, cleaned with 1X Ampure XP beads, resuspended in a final volume of 40ul nuclease-free water and quantified by Qubit. Cut and uncut template was visualized using an Agilent Bioanalyzer high sensitivity DNA chip to confirm 100% digestion. Digested product (0.06 pmol) was ligated into pre-EcoR I/Hind III digested T7 phage vector arms (EMD Millipore, Inc) with 1ul T4 DNA ligase (NEB), 0.5 ul T4 DNA ligase buffer, 1ul of pre-digested T7 vector arms (0.02 pmol) and nuclease-free water to a final reaction volume of 5ul. Four (4) ligations were set up in parallel, incubated at 16C x 12H in a thermal cycler and heat inactivated for 10” at 65C. The entire ligation reaction (5ul) was mixed with 25ul of T7 phage packaging extract (EMD Millipore, Inc) and incubated at room temperature for 2H. After incubation, the packaging reaction was quenched with 120ul of cold Luria-Bertani (LB) broth with carbenicillin added to a final concentration of 1ug/ul. The four packaging reactions were pooled and used to inoculate 1L of *Escherichia coli* strain BLT-5403 grown to an OD_600_ of 0.9 and incubated until the culture lysed (OD_600_ < 0.1), approximately 2.5H. After lysis, 0.1 volumes of 5M NaCl was added to the lysate, it was divided between 500ml polycarbonate centrifugation bottles (Beckman-Coulter) and spun at 12,000 RPM for 30” at 4C in a Sorvall R5-4C centrifuge. The lysate was subsequently filtered using a 0.2 micron bottle-top filter unit (Nalgene, Inc) and 5X PEG-NaCl solution added to a final concentration of 1X. The lysate/PEG mixture was mixed, incubated in a 4C cold room for 12H, centrifuged at 12,000 RPM in a Sorvall R54C centrifuge for 30” at 4C. Pellets were resuspended in Sodium-Magnesum (SM) buffer and plaque forming units quantified by plaque assay as previously described. Resuspended library was prepared for sequencing as described below and paired-end 150 base pair sequencing performed on an Illumina Miseq using a Miseq v2 300 cycle kit.

### Non-human primate infections and samples

Infection of rhesus macaques and collection of samples has previously been described. Serum samples were shipped from WRAIR to San Francisco on dry ice, thawed and randomly aliquoted into plates so that samples from the same individual were not contiguous in the plate. All serum samples were diluted 1:1 with 2X sample storage buffer (40% glycerol, 40mM HEPES pH 7.3, 0.04% NaN3, PBS) and stored in a 4C cold-room after thawing. Samples from mosquito-infected and subcutaneously infected subjects were distributed between two plates.

### Phage immunoprecipitation sequencing

All immunoprecipitation experiments were conducted in 2 ml deep-well polypropylene plates (Genesee Scientific, Inc). Plates were blocked with 3% BSA in TBST overnight with overhead rotation on a Rotator Genie prior to use for immunoprecipitations. After removal of blocking buffer, 1ml of phage library (2 x 10^10^ pfu) was decanted with a multichannel pipet by hand. Serum samples were added directly to each well using an Integra ViaFlow 96 multichannel pipettor, 1ul per well. After addition of serum, plates were sealed with a rubber sealing mat (Genesee Scientific, Inc) and rotated overhead overnight on a Rotator Genie. Plates were removed from the rotator the following morning and spun at 1500RPM in an Eppendorf 5810R centrifuge for 5”. The sealing mat was carefully removed, and 40ul of a mix of Protein A (20ul) and Protein G (20ul) Dynabeads (Thermo-Fisher, Inc) resuspended in Tris/NP-40 was added to each well by multichannel pipet by hand. Protein A and Protein G beads were washed 3x with overhead rotation x 5” in TNP-40 prior to addition to phage-library/serum incubations. Immunoprecipitations were carried out by overhead rotation for 1H at 4C, after which plates were spun at 400 RCF x 1” at 4C in the 5810R centrifuge. The sealing mat was removed, the plate placed on a magnet compatible with 96-well plates and the beads allowed to collect on the magnet and the liquid aspirated. 900ul of RIPA buffer was added to each well, the sealing mat replaced and plates rotated overhead at 4C for 5 minutes. After rotation, plates were spun at 400RCF in the 5810R centrifuge, the sealing mat removed, beads collected on the magnet, wash buffer aspirated and a fresh 900ul of RIPA added. This process was repeated for a total of 5 washes. After the fifth wash, 50ul of phage elution buffer was added to the beads, gently agitated by hand and incubated for 3” on ice. After incubation, beads were collected and inoculated into 1ml of OD_600_ 0.5 *E coli* strain BLT-5403 in a fresh deep-well plate, sealed with a gas-permeable plate sealer and incubated with agitation at 37C at 750rpm using an Infors shaking incubator until all wells showed 100% lysis (1.5-2H). After lysis, 100ul of 5M NaCl was added to each well, mixed and spun at 3220 RPM for 10”. 500ul of spun lysate was added to each well of a blocked deep 96-well plate and an additional 600ul of LB-Carbenicillin added to each well, followed by 1ul of each serum sample to each well. Plates were sealed with a fresh sealing mat and incubated overnight with rotation as already described. The next day, immunoprecipitations and washes were carried out as already described, with the exception that the lysate was not used for an additional round of selection but for sequencing library preparation.

Lysates were heated to 70C for 15 minutes to liberate phage genomes. For each immunoprecipitation, 2ul of lysate was used as the template for a PCR reaction to add Illumina P5 and P7 sequences. Forward and reverse primers for the PCR reaction were AATGATACGGCGACCACCGAGATCTACACNNNNNNNNATGGGCCACGGTGGTCTT CGCCC and CAAGCAGAAGACGGCATACGAGATNNNNNNNNGGGTTAACTAGTTACTCGAGTGC GGCCG, with 96-different barcodes. PCR reactions were done in 25ul volumes with 2ul lysate, 1ul of each primer at 10uM, 0.5ul of 10mM dNTP, 5ul of 5X Phusion buffer, 0.25ul pf Phusion Taq DNA polymerase and nuclease free water to the final volume. PCR conditions were identical to those already listed for library cloning. After PCR, 5ul of each reaction was pooled, cleaned up as already described with Ampure XP beads and quantified using the KAPA Biosystems Illumina quantification kit. All samples were subsequently submitted to the Chan-Zuckerberg Biohub sequencing facility and sequenced on a NextSeq 550 high output (400 million reads) cartridge using custom read 1 and indexing primers. A single-ended 150 base pair sequencing strategy (sequencing 150/186 base pairs of inserts) was used, with a goal read-depth of 1.5-2 millions reads/sample.

### Bioinformatic analysis of PhIP-Seq results

Demultiplexed sequencing data was aligned to the library oligonucleotide sequences using the Bowtie2 aligner. All alignments that were not full-length, perfect alignments were discarded. Perfect alignments were translated and only those that were perfect matches to the input library retained. This set of sequences was tabulated for each of the immunoprecipitation reactions, resulting in a read-count per peptide. Peptide read-counts were converted to reads per 100,000 reads (rp100k) for each sample by dividing by the total number of reads in a sample and multiplying by 100,000.

All samples were organized by animal in temporal order, starting with baseline-pre-infection samples. To determine which hits were significant, we calculated enrichment scores for all identified peptides relative a reference sample for each animal. For samples that were 7, 35, 90 and 365 days post-primary infection, the reference sample was the baseline (pre-infection) samples for each animal. For secondary infection samples, the reference sample was the 15 month pre-secondary infection, post-primary infection sample for each animal. For any peptide not present in the baseline sample, the median rp100k from the reference sample was used to calculate enrichments.

The fit of several probability distributions with enrichment data was examined, including the normal, negative binomial and Poisson. The normal distribution was found to have the best fit, particularly after log transformation of enrichments. In brief, a set of enrichments for a sample was log-transformed and a z-score calculated for each peptide. Peptides with a z-score > 3 were considered as significantly enriched compared to the reference sample.

All peptides in the library were aligned against reference genomes for all four DENV serotypes using blastp. These data were then used to classify peptides as either DENV1 type-specific or cross-reactive. DENV1 type-specific peptides were those that were >= 90% homologous across their full length to DENV1 alone, while cross-reactive peptides were those that were at least 50% homologous to at least 2 DENV strains. Using these data, the proportion of significantly enriched peptides for each timepoint for each animal by each route of infection was determined.

For generating peptide coverage maps, a different strategy was used. Some regions of the flavivirus genome are highly conserved across multiple viruses (e.g. the fusion loop sequence of the envelope protein) and linear peptide sequences recognized by antibodies, such as those represented in the phage display library used here are short. To account for both of these factors, all peptides in the library were decomposed into a series of overlapping 10mers, with each 10mer aligned to the DENV1 reference genome using blastp and only those that were >= 90% homologous to the DENV1 reference genome retained for downstream analysis. For each sample, each significant peptide was mapped to its set of DENV1 homologous 10mers. Using these alignment data and enrichment scores, a cumulative enrichment score was calculated at each position of the DENV1 proteome. As before, these data were computed for each animal, at each timepoint by each route of infection. The median cumulative enrichment art each position was computed and used to generate coverage plots for each timepoint by each route of infection.

### Statistical analysis of phage immunoprecipitation sequencing results

Area under the curve (AUC) was calculated using the median cumulative enrichment data generated as described above. These were calculated for all 10 amino-acid windows across the DENV1 proteome but also for specific regions of interest in the E, NS1, NS3 and NS5 proteins. Permutation testing was used to test for the statistical significance of the differences of AUCs in the mosquito-inoculated and subcutaneously inoculated groups. Briefly, all 20 animals were randomly assigned to the mosquito-inoculated or the subcutaneously inoculated group AUCs calculated and the difference between mosquito and subcutaneously groups calculated. This process was repeated for a total of 1000 iterations, generating the null distribution for AUC_msq_-AUC_sc_ across the DENV1 proteome. The measurements at each distribution were used to model the cumulative distribution function of AUC_msq_-AUC_sc_ and to estimate P(*X* > *x*) at that position.

### ED III ELISAs

Envelope protein residues 301-394 were PCR amplified from a gene block of the Entire DENV1 E protein reference sequence ordered from IDT, Inc. An internal Hind III site was removed by PCR and a second PCR performed to add a 5’ EcoR I site, a 3’ FLAG tag (DYKDDDDK) followed by a 3’ Hind III site. The PCR product was digested in a 50ul reaction as previously described and full digestion confirmed by Agilent Bioanalyzer. Digested product was cloned into an expression vector (OriGene pCMV6), sequenced confirmed by Sanger sequencing (Quintara Bio, Inc.) and CaPO_4_ transfected into 50% confluent HEK293 cells. Transfected cells were lysed with RIPA buffer, to which a protease inhibitor cocktail pill was added (Roche, Inc). Lysates were spun at 13.2K RPM in an Eppendorf microcentrifuge at 4C for 15 minutes, supernatants aspirated, aliquoted in 200-400ul aliquots and stored at −80C until use.

Clear polystyrene plates for ELISAs (R & D Biosystems, Inc) were adsorbed with anti-FLAG antibody (Cell Signalling, Inc.) diluted 1:2500, 50ul per well overnight at 4C in a sealed plate. Plates were washed 3 times with PBS + 0.05% Tween-20 (PBST), 200ul of 2% BSA + PBST added to each well and plates incubated at room temperature for 2H. Blocking buffer was removed, sample diluted 1:100 in blocking buffer added to each well, the plate sealed and incubated overnight at 4C. After sample incubation, all wells were washed 4 times with 300ul PBST per well by multichannel pipet, 50ul of anti-human IgG-HRP conjugated detection antibody (Sigma Chemical, #) added, the plates sealed and incubated at room temperature in the dark for 1H. Plates were washed 5x with 300ul PBST per well as already described and 50ul of TMP Substrate/Peroxidase solution added. Reactions were stopped using 2N H_2_SO_4_ and OD_450_ measured on a plate reader. For each sample series, the ratio of the OD_450_ to pre-infection OD_450_ was calculated. Median ratios were compared using a one-sided Mann-Whitney U test to determine the statistical significance of the observed differences.

### Analysis of antibody binding specificities

For each animal at each timepoint by each route of infection, the significantly enriched peptides that spanned residues 347-377 of the E protein were identified. Multiple sequence alignments were performed using the package DECIPHER and the proportion of aligned sequences at that position whose amino acid residue matched that of the reference sequence was calculated.

### Correlation analysis with neutralization data

We used the R function *cor* to calculate Pearson correlation coefficients between log transformed ELISA data and 13 month geometric mean neutralization titers (GMT) against DENV1 that had been determined for each of the 20 animals. For the analysis of correlation between phage data and the GMT data, we calculated the sum of all enrichments for all possible 45 amino-acid windows for each animal for both the mosquito and subcutaneously inoculated groups. We then correlated these vectors of summed enrichments with GMTs and plotted the resulting correlation coefficients for each position. All of these analyses were conducted using R.

### Disclaimer

Material has been reviewed by the Walter Reed Army Institute of Research. There is no objection to its presentation and/or publication. The opinions or assertions contained herein are the private views of the author, and are not to be construed as official, or as reflecting true views of the Department of the Army or the Department of Defense. Research was conducted under an approved animal use protocol in an AAALACi accredited facility in compliance with the Animal Welfare Act and other federal statutes and regulations relating to animals and experiments involving animals and adheres to principles stated in the *Guide for the Care and Use of Laboratory Animals*, NRC Publication, 2011 edition.

## ACKNOWLEDGEMENTS

This work was supported by Department of Defense grant D16AP0002 to JD (PI).

