## Supplemental Figures for "Inoculation by mosquito induces durable differences in serological profile in non-human primates infected with DENV1"

### Supplemental Data

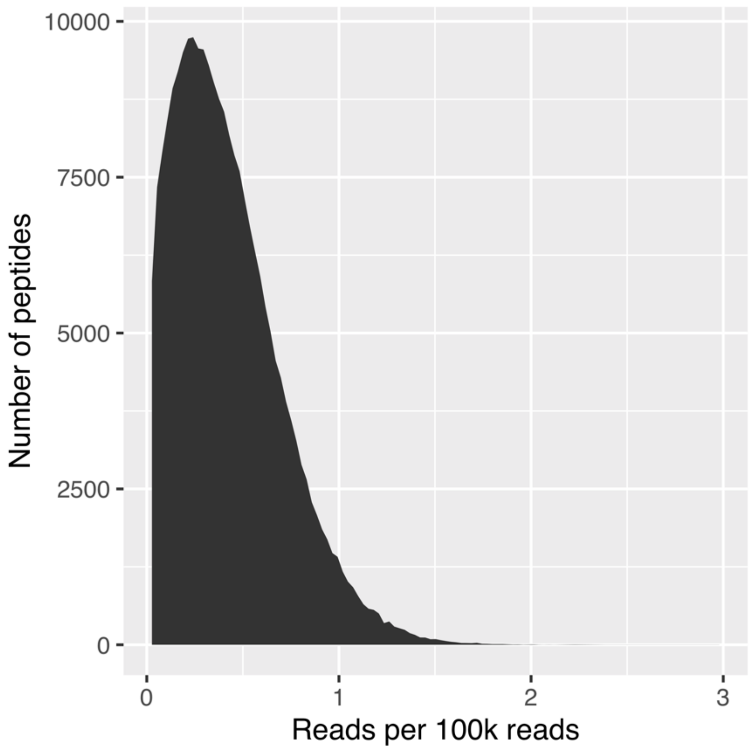


**Supplemental Figure 1. Peptide representation of packaged phage display library**. Library oligonucleotides were amplified and cloned into the T7 phage display vector as described in Methods. At a sequencing depth of 10.7 million reads, 96.7% of designed peptides were represented. Of the peptides represented, 96% were present at less than or equal to 1 read per 100,000 reads. The x-axis is log scaled.

A

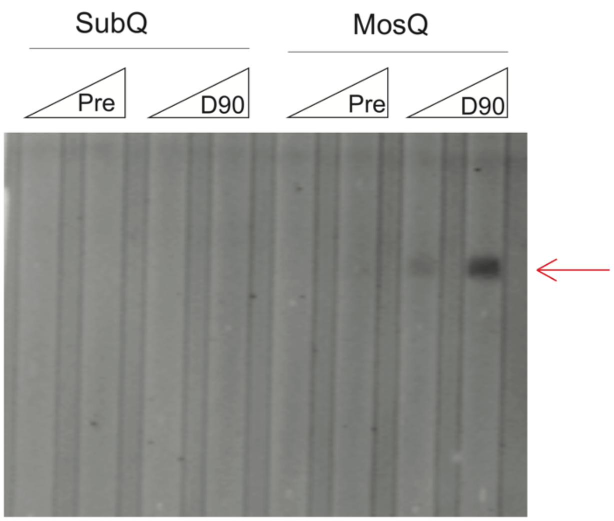


B

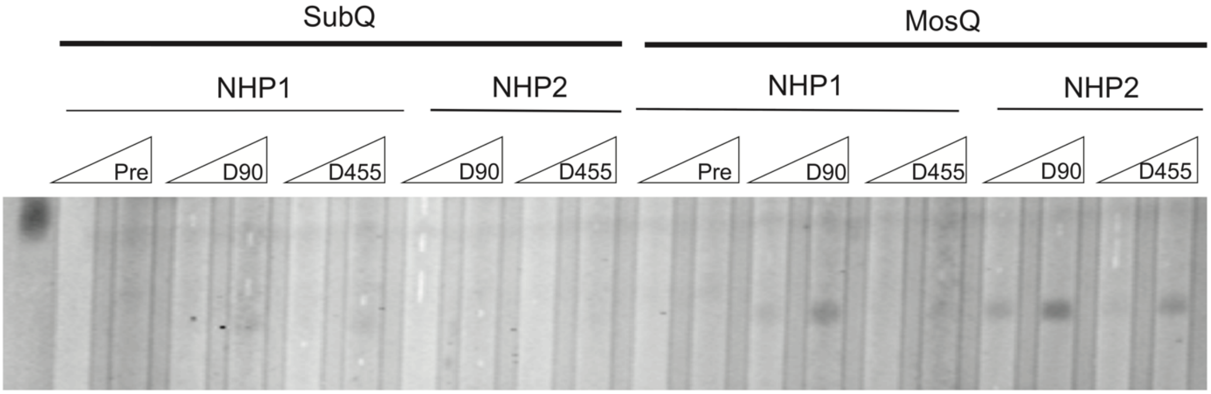


**Supplemental Figure 2. Persistent antibodies against envelope protein residues 339-384 in macaques inoculated by mosquito.** Slot immunoblots were performed using sera from macaques inoculated by mosquito bite and by subcutaneous injection. Results in 1 pair of animals showed that the signal was absent at baseline in both groups but present at 90 days post-infection, with the signal titrating (A). The same results were obtained in two other animals from each group, extending to 455 days post-infection (B). These results were consistent with PhIP-Seq results. All Sera were diluted 1:100 in PBS for immunoblots.

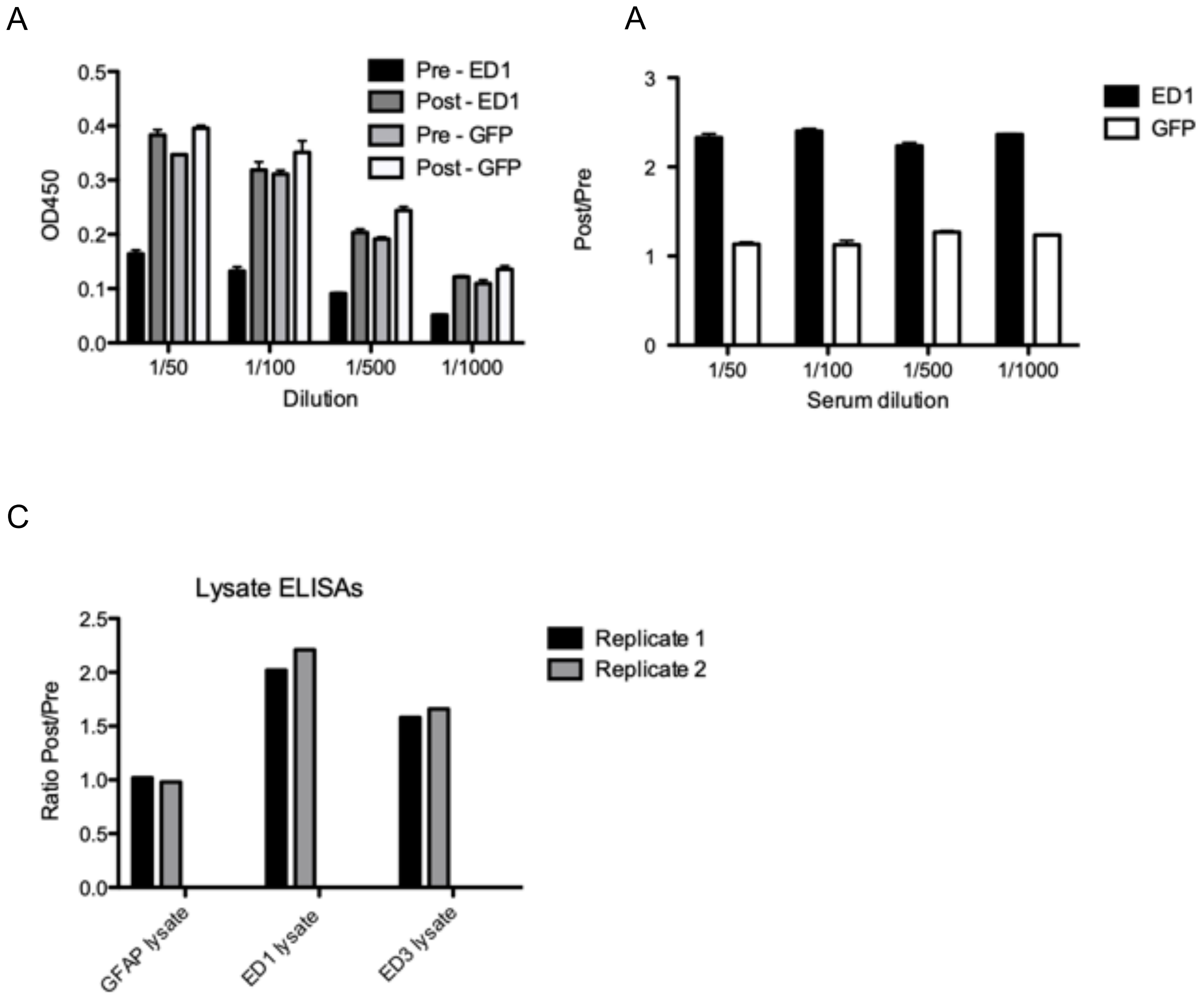


**Supplemental Figure 3. Titrations of ELISAs for two different envelope protein domains that are enriched by PhIP-Seq.** Constructs spanning envelope protein residues 1-91 and 339-384 were cloned into a mammalian expression vector and overexpressed in HEK293T cells. A FLAG tag was added to the carboxy terminus of each construct. Whole cell lysates were captured using an anti-FLAG antibody (Cell Signaling, Inc), diluted macaque sera added and detected with a an anti-monkey IgG HRP conjugated antibody (Sigma, Inc). Signal, as measured by OD450 for both constructs titrated with decreasing amounts of serum (A). The same was true for OD450 ratios of post-exposure (90 days) to pre-exposure (B). Ratios for a control construct (GFP) remained ~1.0, while those for a construct against ED1 and ED3 increased 90 days post-exposure (C).
